# FARCI: Fast and Robust Connectome Inference

**DOI:** 10.1101/2020.10.07.330175

**Authors:** Saber Meamardoost, Mahasweta Bhattacharya, EunJung Hwang, Takaki Komiyama, Claudia Mewes, Linbing Wang, Ying Zhang, Rudiyanto Gunawan

## Abstract

The inference of neuronal connectome from large-scale neuronal activity recordings, such as two-photon Calcium imaging, represents an active area of research in computational neuroscience. In this work, we developed FARCI (Fast and Robust Connectome Inference), a MATLAB package for neuronal connectome inference from high-dimensional two-photon Calcium fluorescence data. We employed partial correlations as a measure of the functional association strength between pairs of neurons to reconstruct a neuronal connectome. We demonstrated using gold standard datasets from the Neural Connectomics Challenge (NCC) that FARCI provides an accurate connectome and its performance is robust to network sizes, missing neurons, and noise levels. Moreover, FARCI is computationally efficient and highly scalable to large networks. In comparison to the best performing algorithm in the NCC, FARCI produces more accurate networks over different network sizes and subsampling, while providing over two orders of magnitude faster computational speed.

## Introduction

The human brain comprises about 100 billion neurons, each making thousands of synaptic connections with others [1]. Neuronal connectome, the wiring of neurons, is highly plastic and dynamic. The plasticity of the connectome is a feature that imparts the brain with an ability to learn new behavior and to store and process new information [2]. The inference of brain’s neuronal connectivity has received much attention and is one of the main scientific challenges in the 21^st^ century. For example, the Human Connectome Project (HCP) [3,4] was established in 2009 with the goal of mapping the human’s brain connectome. Knowledge of neuronal connectome and its rewiring may shed light on the operating principles of the brain and its myriad functions.

Direct measurements of the connectome based on anatomical techniques is time-consuming, non-scalable and challenging due to limitations of macroscale imaging modalities [5]. Therefore, an alternative approach is to indirectly infer connectome based on neuronal activity recording data. Technological advances in neuroscience have enabled the recording of neuronal activity signals in awake animals at high resolution [6], accelerating efforts toward neuronal connectome inference [1,7]. For instance, two-photon Calcium imaging and neuronal electrophysiology are two current technologies for measuring the activity of single neurons. While electrophysiology offers higher temporal resolution which enables recording the activity of fast spiking neurons, Calcium imaging provides a high spatial resolution as it can record a large population of neurons that are adjacent to one another [8]. Although these techniques employ different principles to detect neuronal activity signals, both technologies are capable of generating large scale time series data for 100s-1000s neurons simultaneously.

Neuronal activities are the outputs of the neuronal networks, and do not directly indicate the connections among the neurons. For this reason, state-of-the-art computational algorithms have been developed to reconstruct the neuronal connectome from high throughput time-series neuronal firing data. In general, there exist two classes of methods for inferring neuronal connectivity from neuronal activity measurements: model-free methods and model-based methods (see [7] for a more comprehensive review). Each class of methods captures the neuronal connectome with different details such as strength, type (i.e. excitatory or inhibitory), and directionality of the connections. Model-free methods make no assumption on the biological mechanism that generates the observed data, using statistical inference, information theory, and supervised learning approaches, to reconstruct the connectome [7]. Model-based methods are typically more complex by employing a generative model of the neuronal network activity. Here, the connectivity is inferred by explicitly modeling the activity of pre-and post-synaptic neurons and the connectome inference is formulated as an estimation of some model parameters so that the model is capable of explaining the observed neuronal activity patterns. Models generated from such model-based methods can further be used to generate neuronal activity data for validation and prediction. But, in practice, it is not feasible to account for all aspects governing the observed neuronal activity data (e.g. response to external stimuli, synaptic strength and connectome rewiring) in a single model. Consequently, model-based approaches use simplified models with the aim of only simulating empirical data without guaranteeing an explicit correspondence between model parameters and physiological factors [9]. Results of the Neural Connectomics Challenge (NCC) [10,11] revealed that simpler model-free methods, especially those relying on partial correlation coefficients, outperform more complex methods [12]. Note that the partial correlations are symmetric and thus do not give an indication for the directionality of the neuronal connections. Nevertheless, an undirected neuronal connectome inferred from neuronal activity recordings can facilitate the comprehension of functional neuronal circuits and potentially of their rewiring principles during learning and other brain functions.

In this work, we developed a model-free method FARCI (Fast and Robust Connectome Inference), a MATLAB toolbox for inferring neuronal connectome from time-series Calcium fluorescence data of neuronal activity. FARCI combines non-negative deconvolution, thresholding, and smoothing in data pre-processing and produces a partial correlation network for neuronal connectome reconstruction. We assessed the performance of FARCI using gold standard Calcium fluorescence datasets from the NCC [12]. Our results demonstrate that FARCI is highly efficient and scalable to large dataset and connectome, and its performance is superior to the winner of the NCC in terms of accuracy and robustness to noise levels, sampling rates, network densities, and hidden (missing) neurons.

## Methods

### Spike deconvolution

Precise temporal information of individual neurons’ spiking activity is crucial for connectome inference. Two-photon Calcium (Ca^2+^) imaging indirectly captures neuronal action potentials via fluorescence signals of a genetically encoded Calcium indicator (GECI). Extracting neuronal spiking activity from Ca fluorescence data is a non-trivial task because raw Ca imaging recordings are known to have significant intrinsic noise, baseline fluorescence drift, and other technical constraints, such as low sampling rate and slow decay of fluorescence sensor [13]. Among various algorithms for the inference of neuronal spiking activity from two-photon imaging data, Non-Negative Deconvolution (NND) method has been shown to outperform other algorithms while having simple parameter settings [13,14].

FARCI uses a sparse NND, using the Online Active Set method to Infer Spikes (OASIS) algorithm [15] from the Suite2P MATLAB package [16]. OASIS is an adaptation of the pool adjacent violators algorithm [17]. Here, we write the neuronal spike deconvolution, as follows:

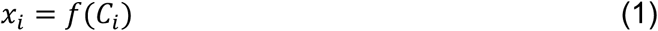

where *C* is the raw Ca fluorescence data of neuron *i, f* represents the deconvolution function, and *x* is the deconvolved neuronal spiking activity.

### Spike denoising

The deconvolved spiking activity are often contaminated by noise and such noise can degrade the accuracy of the inferred connectome. For this reason, we remove the deconvolved neuronal spike *x* that is below a certain neuron-specific threshold. The neuron specific threshold Θ_*i*_ is set to the average *x* of neuron *i* plus a user-specified constant multiple of the standard deviation. The denoised spiking activity *y*_*i*_ is computed as follows:

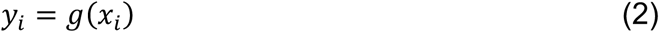

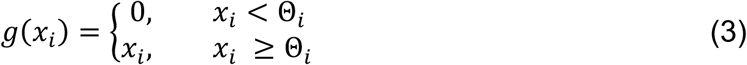

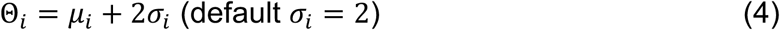

where Θ_*i*_ is the denoising threshold and *μ*_*i*_ and *σ*_*i*_ are the mean and standard deviation of the spiking activity for neuron *i* – denoted by *x*_*i*_, respectively. While denoising may remove actual spiking activity with a low amplitude, our tests below indicate that the accuracy gain by spike denoising outweighs the loss of information due to removal of low-amplitude spiking activity.

### Spike smoothing

Binning spikes over multiple time points (image frames) has been shown to enhance the correlation between deconvolved and ground-truth spiking activity [18]. We tested different weighted binning strategies for smoothing the spike data (see Supplementary Table S1). We identified the following smoothing function *h*(*y* _*i*_, *t*) as the best among the strategies:

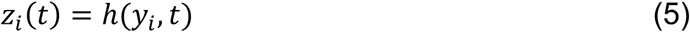

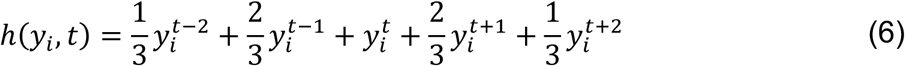

where *h* is the smoothing function, *z*_*i*_ denotes the smoothed spikes, and *t* denotes the time index of spiking activity. The data pre-processing pipeline from Ca fluorescence data to the final neuronal spikes is illustrated in Figure 1.

**Figure 1.**
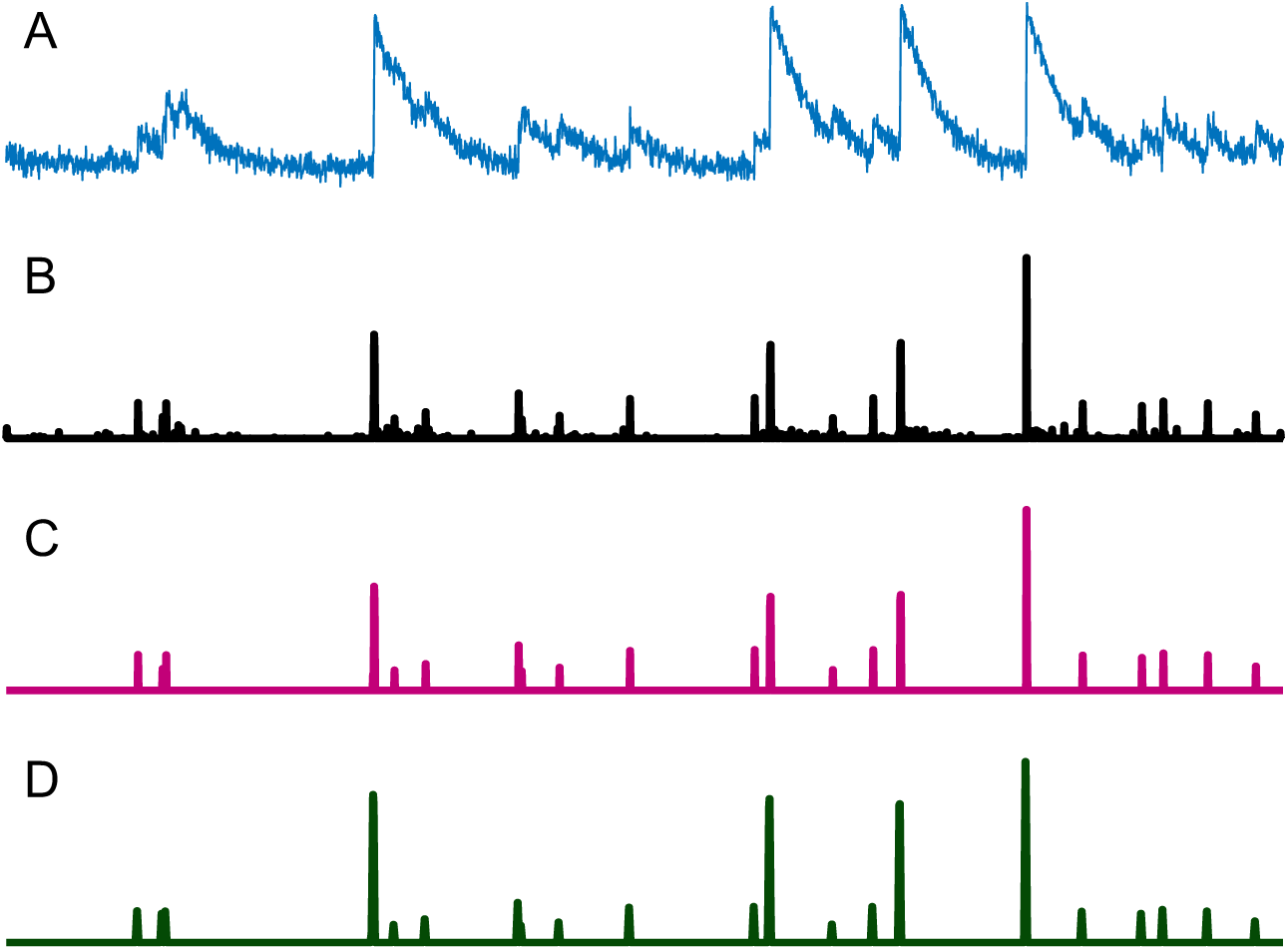
Data preprocessing pipeline of Calcium fluorescence data. **A**. Raw Ca fluorescence signal. **B**. Deconvolved spikes using OASIS. **C**. Denoised spikes. **D**. Smoothed spikes.

### Partial correlation statistics

In FARCI, the functional connectivity between each pair of neurons are established based on their co-firing behavior. However, due to highly interconnected nature of neurons, functional correlations between any two neurons may arise indirectly from their connections to other neurons (e.g., sharing the same pre-synaptic neurons). To reduce false positives, FARCI uses the partial correlation coefficient as a measure of functional connectivity between a pair of neurons – that is, the correlation between the spike activity of two neurons while controlling for the activity of other neurons. In order to infer an edge (connection) between two neurons, for a set *V* of *N* neurons, we treat the spike data of the *N* neurons at each time point as *N* independent and identically distributed (*i. i. d*.) observations, i.e. *Z* = {*z*;_*1*_, *z*_*2*_, …, *z*_*n*_ }. The partial correlation *p*_*ij*_ between neuron *i* and *j*, is computed via the precision matrix ϕ as follows:

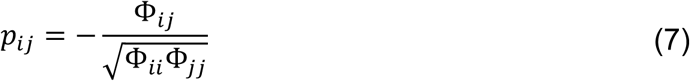

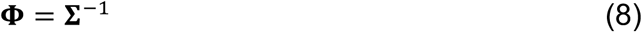

where **Σ** is the *N* × *N* covariance matrix of the neuronal spikes *z* and **Φ** is the precision matrix – the inverse of the variance-covariance matrix. The partial correlation coefficients have values between –1 and +1, where a value of +1 (–1) indicates a perfect positive (negative) correlation of spike activity between two neurons when all other neurons having their activity fixed.

### Performance evaluation

We evaluated the performance of FARCI using the gold standard *in silico* Ca fluorescence data and connectome in the NCC. The provided datasets in NCC were simulated using a realistic and state-of-the-art simulator [19], generating data that resemble actual recording of Calcium signals from neurons. The data simulator employed a mathematical model that takes into account limitations of Calcium imaging technology such as temporal resolution and light scattering artifacts. We followed the assessment of method performance in the NCC and evaluated the area under Receiver Operating Characteristic (AUROC) curve by comparing the inferred partial correlation network with the ground truth connectome. Besides, we use the area under Precision-Recall (AUPR) curve as an additional performance metric. AUROC and AUPR values range between 0 and 1, where a value of 1 indicates a perfect prediction. Note also that an AUROC of 0.5 is the expected performance for a random prediction. Although AUROC and AUPR are related to each other, unlike AUROC, AUPR takes into account the ratio between positives and negatives (i.e. class imbalance). For a sparse connectome where the number of true connections is low in comparison to the number of all possible connections, AUPR is a more sensitive measure for the performance of network inference methods than AUROC [20]. On the other hand, AUROC tends to be very high (near 1) for a sparse network.

We compared FARCI with the best performer in the NCC [12], an algorithm developed by Sutera et al., 2015. The Sutera et al. algorithm comprises a four-step signal processing pipeline (low-pass filter, high-pass filter, hard thresholding, and weighting), and like FARCI, produces partial correlation networks. We noted that the Sutera et al. algorithm gave an excellent performance for 1000 neuron connectomes in the NCC – the connectomes used for scoring in the challenge. However, its performance was relatively poorer for the smaller connectomes in the NCC with 100 neurons. In other words, the Sutera et al. algorithm appeared to be overly optimized, and thus is not robust to network size. In this work, we also evaluated the robustness of FARCI performance with respect to not just network sizes, but also noise level, sampling rate, and hidden (missing) neurons.

## Results and Discussion

FARCI is an efficacious and robust method for inferring functional neuronal connectome from Calcium fluorescence data. Figure 2 illustrates the workflow of the connectome inference in FARCI, which comprises the following key steps: (1) deconvolution of spiking activity from Ca fluorescence data, (2) spike denoising, (3) spike smoothing, and (4) evaluation of partial correlations. The functional neuronal connectome is represented by the partial correlation network among the neurons in the dataset. We benchmarked FARCI using gold standard Ca fluorescence datasets from the Neural Connectomics Challenge (NCC) [10,11]. The NCC provides a total of 16 gold standard datasets generated by *in silico* simulations [19] of the activity of 100 (*n =* 6) and 1000 neurons (*n* = 4) that are referred to as the *small* and *normal* connectomes, respectively. Additional datasets of 1000 neurons (*n* = 6) with different characteristics of noise, neuronal firing rate, network clustering coefficient, and network density are also available. More details of the datasets in the NCC are provided in Table 1. The connectivity of the small and normal connectomes has a relatively low density that ranges within 16.3 ± 1.7% for the small networks and 2.1 ± 0.5% for the normal networks, suggesting that these connectomes are sparse.

**Table 1.** Datasets provided in Neural Connectomics Challenge. Each dataset contains three types of information: 1. neuronal activity in the form of Ca fluorescence signals, 2. the ground truth connectome structure, and 3. the spatial coordinates of neurons.

| Networks | # of neurons | Description |
| --- | --- | --- |
| small (6 datasets) | 100 | six connectomes with 100 neurons |
| normal (4 datasets) | 1000 | four connectomes with 1000 neurons |
| normal3-highrate | 1000 | normal-3 connectome with highly active neurons |
| normal4-lownoise | 1000 | normal-4 connectome with a higher signal to noise ratio |
| highcc | 1000 | a connectome of 1000 neurons with a higher clustering coefficient |
| lowcc | 1000 | a connectome of 1000 neurons with a lower clustering coefficient |
| highcon | 1000 | a connectome of 1000 neurons with a higher number of connections per neuron |
| lowcon | 1000 | a connectome of 1000 neurons with a lower number of connections per neuron |

**Figure 2.**
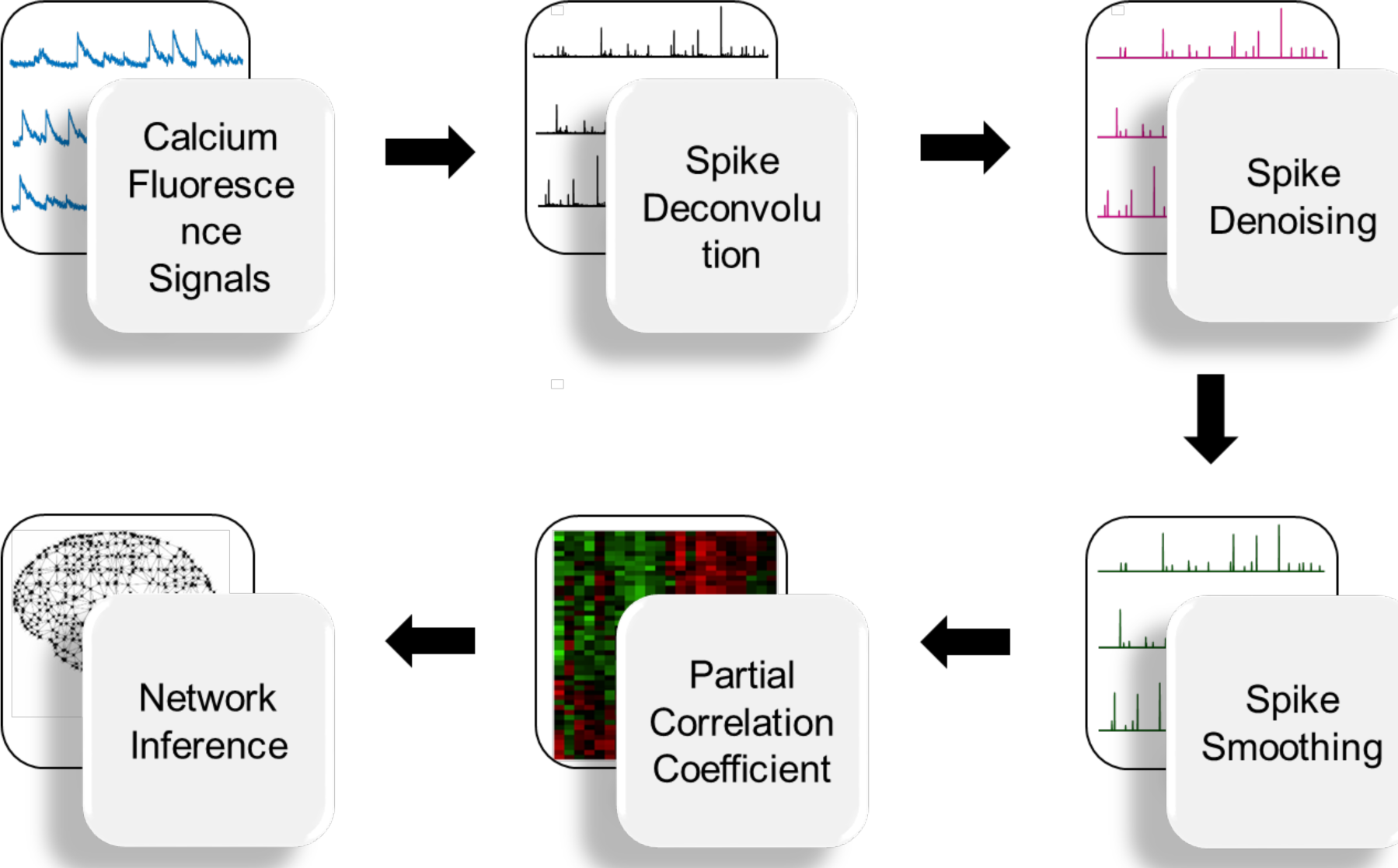
Workflow of connectome inference in FARCI. FARCI combines denoising and smoothing of neuronal spikes, the output of which is used to generate partial correlation networks.

In the NCC, participants were asked to reconstruct neuronal connectome from time-series measurements of Ca fluorescence data. Submissions from the participants were ranked based on how accurately their algorithms are able to infer the neural connectomes with 1000 neurons – the so-called normal connectome. The scoring in the NCC was done by evaluating the area under the receiver operating characteristic (AUROC). But, as we noted in Methods, AUROC tends to be high for a sparse connectome, and hence, is not a sensitive measure of performance. For this reason, in our benchmarking, we evaluated the AUROC as well as the area under the precision-recall curve (AUPR) (see Methods) to assess the performance of FARCI. The AUPR is a more sensitive indicator for network inference accuracy than the AUROC for a sparse connectome [20]. Below, we compared FARCI with the best performing method in the NCC by Sutera *et al*. [12] in terms of AUROC and AUPR.

### Neuronal spike deconvolution

Ca fluorescence imaging data give only indirect measurements of neuronal activity, and thus, require data pre-processing to extract the underlying neuronal action potential spikes. We employed OASIS deconvolution algorithm [15] from the MATLAB package Suite2P [16] that uses a non-negative deconvolution strategy to provide estimates for timing and amplitude of spiking activity. Table 2 gives the AUROC and AUPR values for using the partial correlations of the deconvolved spikes to infer neuronal connectomes (see Table S2-S4 for more detailed results). While the AUROCs were generally good (>0.78), the AUPRs were as low as 0.23.

**Table 2.** Effect of different signal processing steps on connectome inference. Unlike binarization, signal thresholding and smoothing improve the accuracy of connectome inference.

| Filter | AUROC |  |  | AUPR |  |  |
| --- | --- | --- | --- | --- | --- | --- |
|  | small<br>(n = 6) | normal<br>(n = 4) | others<br>(n = 6) | small<br>(n = 6) | normal<br>(n = 4) | others<br>(n = 6) |
| Neuronal<br>Spikes<br>$x$ | $0.871 \pm 0.058$ | $0.892 \pm 0.002$ | $0.905 \pm 0.030$ | $0.543 \pm 0.097$ | $0.335 \pm 0.005$ | $0.347 \pm 0.067$ |
| Spikes<br>+ Binarization<br>$u(x)$ | $0.563 \pm 0.055$ | $0.653 \pm 0.006$ | $0.647 \pm 0.069$ | $0.189 \pm 0.021$ | $0.042 \pm 0.002$ | $0.050 \pm 0.021$ |
| Spikes<br>+ Thresholding<br>$g(x)$ | $0.876 \pm 0.051$ | $0.882 \pm 0.003$ | $0.895 \pm 0.040$ | $0.620 \pm 0.096$ | $0.408 \pm 0.012$ | $0.421 \pm 0.114$ |
| Spikes<br>+ Smoothing<br>$h(x)$ | $0.891 \pm 0.035$ | $0.908 \pm 0.002$ | $0.917 \pm 0.025$ | $0.538 \pm 0.056$ | $0.330 \pm 0.004$ | $0.346 \pm 0.053$ |
| **FARCI**<br>$h(g(x))$ | **$0.916 \pm 0.031$** | **$0.908 \pm 0.002$** | **$0.918 \pm 0.032$** | **$0.741 \pm 0.067$** | **$0.491 \pm 0.003$** | **$0.497 \pm 0.100$** |

### Binarization of neuronal spikes

We also tested whether binarizing the deconvolved spiking activity might help in improving connectome inference using partial correlations. As reported in Table 2, converting spikes to binary data – by setting all non-zero spikes to 1 – led to a significant deterioration in the accuracy of the inferred connectomes for both small and normal sized networks. The result above suggests that the amplitude of spiking activity contain significant information for inferring neuronal connectivity. Thus, in FARCI, we used the deconvolved spiking activity without any binarization.

### Neuronal spike denoising

Low amplitude spiking activity may arise from random noise, and should ideally be removed to improve accuracy. In FARCI, we implemented a denoising step by enforcing a user defined threshold that is equal to a multiple *α* of the standard deviation of Ca spike heights. More specifically, we define the threshold Θ for each neuron *i* as follows:

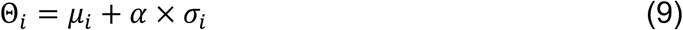

where *μ*_*i*_ and *σ*_*i*_ denote the mean and standard deviation of the deconvolved spiking activity *x*_*i*_. During denoising, any spiking activity below the threshold is set to 0.

We investigated the influence of *α* on AUROC and AUPR. To this end, we ran the connectome inference using denoised spikes for different *α* values in the range of 0 ≤ *α* ≤ 5. As shown in Figure 3, the AUROC generally drops with increasing denoising strength (i.e. increasing *α*), especially for larger connectomes, but stays reasonably high at above 0.7. Indeed, for large and sparse networks where the number of negative cases (i.e. the absence of neuronal connections) significantly outweighs the number of positive cases, AUROC often becomes too optimistic. Here, the AUPR serves as a more sensitive metric for method performance. The AUPRs for all of the connectomes show a peak for *α* between 2 and 3, with *α* = 2 often giving the highest value. For this reason, we use *α* = 2 as the default value for the spike denoising step in FARCI. As reported in Table 2, the denoising step using *α* = 2 achieves an average improvement of 18.6% in the AUPR.

**Figure 3.**
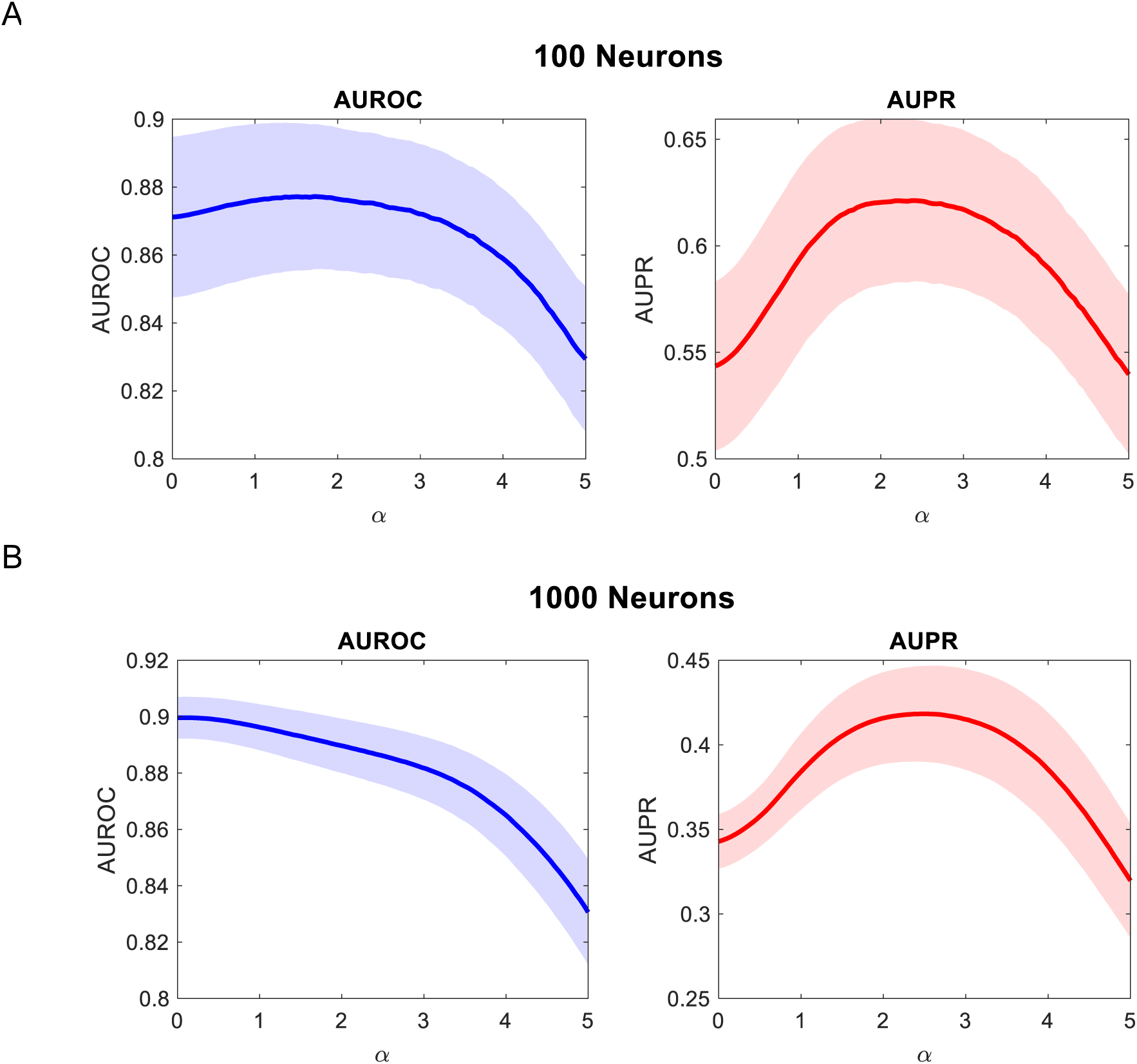
Effect of signal denoising on AUROC and AUPR in networks of **A**. 100 (n = 6), and **B**. 1000 neurons (n = 10). While AUROC tends to drop with increasing *α*, AUPR reaches a peak for values of 2 < *α* < 3. The shaded area denotes the standard error of the mean (SEM).

### Neuronal spike smoothing

Smoothing deconvolved spiking activity has been demonstrated previously to improve the connectome inference [12]. Similarly, binning spikes from OASIS increases the correlation between the predicted and the ground truth spikes [18]. We explored a set of heuristic binning and smoothing functions for improving accuracy (see Supplementary Table S1). Our explorations using various smoothing functions (data not shown) led to Eq. 6 as a simple-yet-efficacious weighted binning function with a superior connectome inference performance for the small and normal connectomes in the NCC. The smoothing function uses a time window of 5 frames where higher weights are given to the time points closer to the center frame of the window. The performance of the connectome inference using binned spiking activity is reported in Table 2, which shows moderate improvements in the AUROC over using the spiking activity directly without binning.

### FARCI

FARCI combines denoising and smoothing of the deconvolved spiking activity to produce a synergistic improvement in the connectome inference, as shown in Table 2. Importantly, the performance comparison in Figure 4 (also see Supplementary Table S5) shows FARCI generally outperforming the best performing algorithm in the NCC by Sutera et al. [12]. The complete results of the benchmarking and comparison are provided in Table S5. The algorithm by Sutera et al. appeared to be optimized for the normal connectomes with 1000 neurons, which is the size of connectomes used for scoring in the NCC. Sutera et al. algorithm performed poorer on the small connectomes with 100 neurons than on the normal connectomes. FARCI is able to outperform Sutera et al. algorithm, providing similarly high AUROC and generally much higher AUPR, regardless of the size of the networks, the level of noise, the density of the networks, and the sampling rates (see Supplementary Table S5).

**Figure 4.**
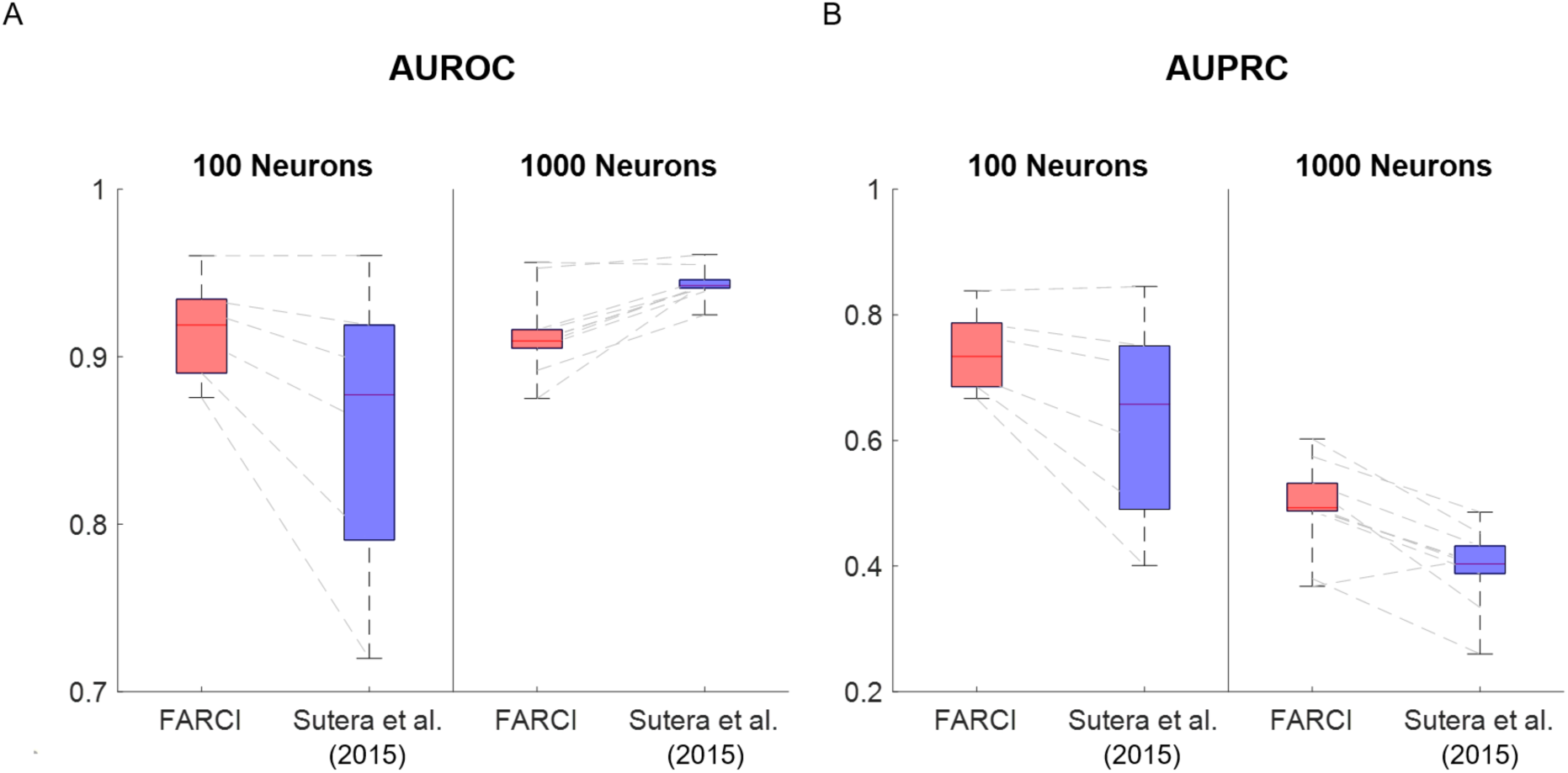
Comparison of FARCI performance with the winner of NCC. The accuracy of the inferred connectome is measured by (A) AUROC and (B) AUPRC.

### Missing neurons

Finally, we investigated the robustness of FARCI with respect to missing neurons. The missing neurons can be considered as hidden variables in the connectome inference. Hidden variables are a common problem in functional connectome inference as only a subset of neurons are measured in a typical experimental setup. We emulated missing neurons by randomly sampling a subset of neurons from the dataset. We then applied FARCI to obtain partial correlation networks for the subsampled neurons, and compared the inferred connectome with the appropriate subnetwork of the ground truth connectome associated with the subsampled neurons. We generated five random samples of neurons and their Ca fluorescence data from the connectomes with 1000 neurons, with the following sizes: 50, 200, 400, 600, and 800 neurons. For each random sample, we performed FARCI and evaluated the average of AUROC and AUPR and the computational runtime. We again compared FARCI with the algorithm by Sutera et al. (2015). Figure 5A-B depicts the AUROC and AUPR from missing neurons simulations using the Normal-1 connectome in the NCC. The results indicate that FARCI is able to maintain high AUROC and AUPR, even up to 60% missing neurons in the dataset. While the AUROC often stays high (>0.9), the AUPR drops quickly at >80% missing neurons. Both FARCI and Sutera et al. (2015) provide comparable AUROCs, but FARCI consistently gives higher AUPR across different fractions of subsampling than Sutera et al. (2015) method. Figure 5C-D summarizes the performance of FARCI for all 1000-neuron networks in the NCC (*n* = 10), confirming the robustness of FARCI to missing neurons up to 60%-80% of the connectome.

**Figure 5.**
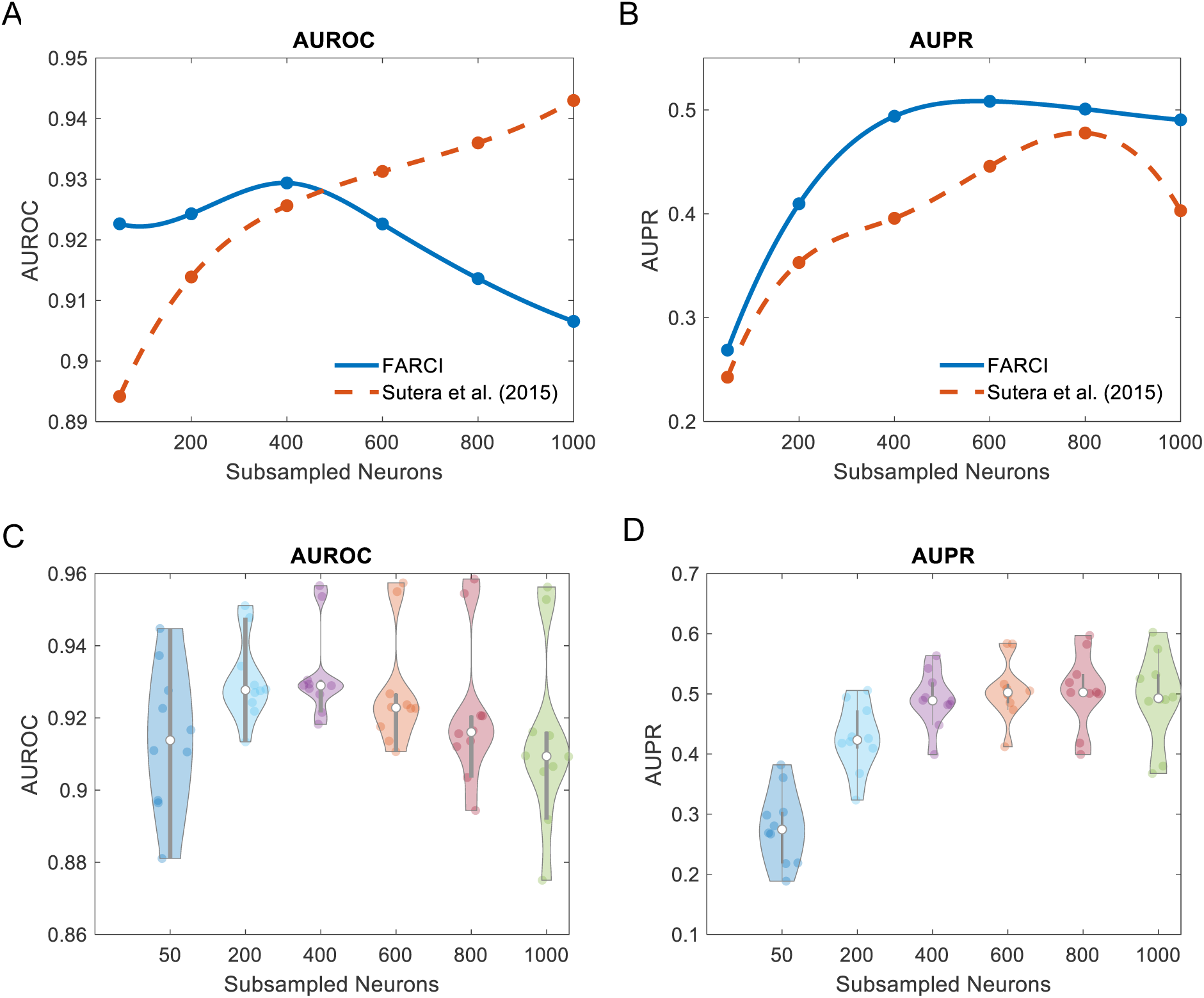
Evaluation of FARCI performance for connectome inference with missing neurons. **A & B**. Comparison of FARCI with Sutera et al. (2015) algorithm in terms of AUROC and AUPR, using the first 1000-neuron connectome from the NCC (Normal-1 network). **C & D**. Performance of FARCI for all 1000-neuron datasets (n = 10).

### Computational speed

Besides accuracy, computational efficiency is a desirable feature of a connectome inference algorithm. As shown in Figure 6A, FARCI offers 2-3 orders of magnitude of computational speed-up over Sutera et al. algorithm over various network sizes. In addition, the computational runtimes of FARCI has a better scaling with network size than Sutera et al. – that is, a lower fold-increase in computational times with increasing connectome size. The fast performance of FARCI is consistently observed across the datasets in the NCC, as shown in Figure 6B. Furthermore, Figure 6B indicates that the runtime of FARCI scales linearly with the size of the connectome, at least up to 1000 neurons.

**Figure 6.**
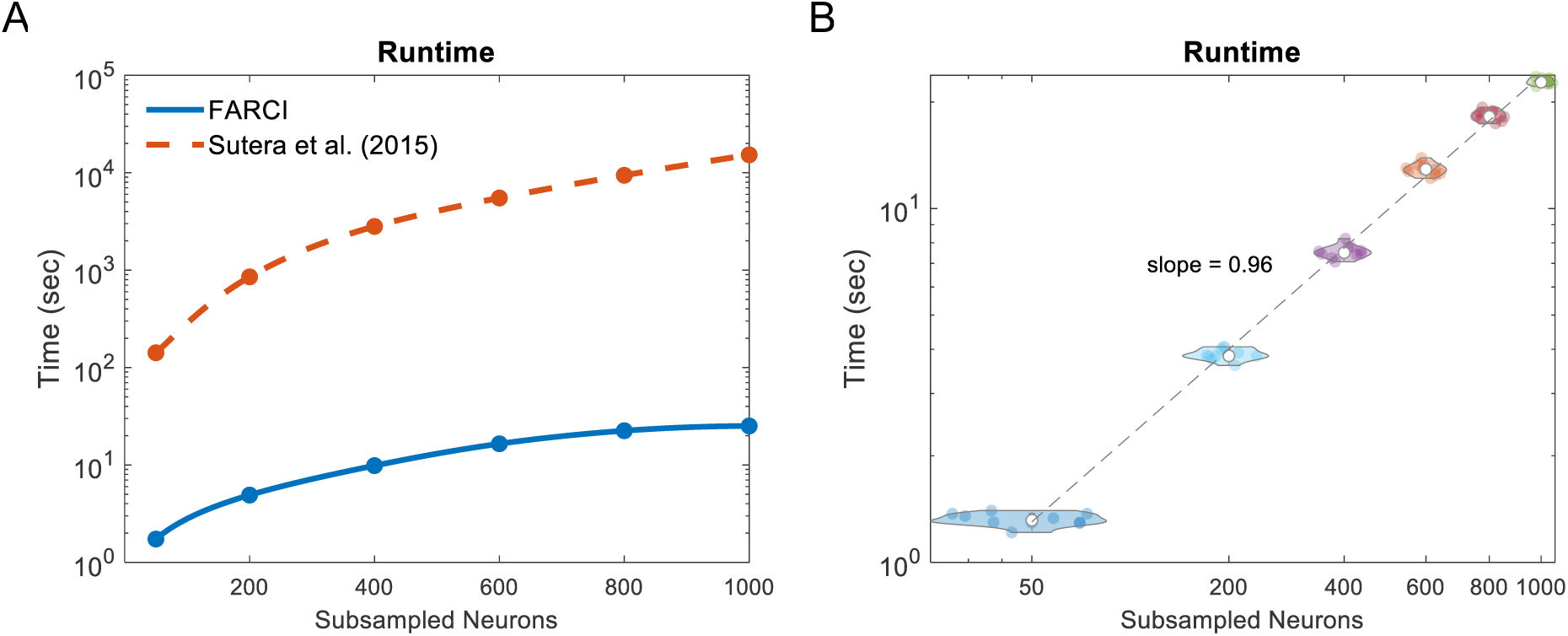
Computational runtimes. **A**. Runtime comparison of FARCI and Sutera et al. (2015) algorithm using Normal-1 network in the NCC. **B**. FARCI runtimes for different sizes of subsampled networks for 1000-neuron datasets from the NCC (*n* = 10).

## Summary

In this work, we developed FARCI, a fast and robust procedure for inferring functional neuronal connectome from two photon Calcium imaging data. FARCI combines a fast non-negative deconvolution algorithm OASIS [15], spike denoising, and spike smoothing, to extract information for neuronal spike events from Ca fluorescence signals. FARCI produces a partial correlation network of the neurons for connectome inference. Benchmarking using gold standard *in silico* datasets from the Neural Connectomics Challenge [10,11], showed that FARCI outperforms the best algorithm in the NCC by Sutera et al. (2015) in terms of the accuracy of the connectome inference (i.e., AUROC and AUPR) and the computational runtimes and scaling. Also, the high performance of FARCI is robust with respect to the connectome size, data noise and sampling rate, and network densities. Finally, we demonstrated that FARCI performs well in the realistic scenario where there are missing neurons in the connectome inference. In this scenario, partial correlations between any two neurons may appear because they shared a pre-synaptic neuron that is not part of the measurement. In our tests, FARCI is still able to maintain high AUROC and AUPR up to 60% missing neurons in a connectome of 1000 neurons, while keeping moderately high AUPR until 90% neurons are missing.

FARCI produces a partial correlation network for the functional connectome, and such networks do not provide indications on the directionality of the neuronal connectivity. The directionality of the neuronal connectivity gives information regarding the identity of the pre- and post-synaptic neurons. However, the challenges for determining directionality in neuronal connectome from two-photon Ca fluorescence data are many. First, the typical rate of data sampling for two-photon Ca imaging ranges between 30 - 100 ms (i.e. ∼10-30 Hz) [21], which is much longer than the time scale of neuronal action potentials and the following refractory period between 1 - 5 ms [22]. Given the sampling rate of Ca imaging, neurons may have fired several times in between any two image frames, and thus the sequential timing of pre- and post-synaptic neuron firing has a low chance to be captured accurately. Other fundamental challenges in determining causal (directional) connectivity from time series data have been discussed elsewhere [7]. Despite the lack of directionality in the connections, the generated functional connectome is still useful for studying connectome dynamics and rewiring in various brain functions, such as in learning and memory formation.

## Availability

The codes, user manual, and tutorial of FARCI are available online (https://github.com/CABSEL/FARCI and http://www.projectmemonet.org/farci).

## Acknowledgement

The authors would like to acknowledge funding support from NSF-HDR IDEAS Lab (funding # 1939987, 1940202, 1940162, 1939999, and 1939992).

## Supplementary Materials

**Table S1.** Smoothing functions. The functions obtained from [12] lead to a poorer neuronal connectome inference performance compared to Eq. 6.

|  |
| --- |
| $f_1(y_i, t) = y_i^{t-1} + y_i^t + y_i^{t+1}$ |
| $f_2(y_i, t) = 0.4y_i^{t-3} + 0.6y_i^{t-2} + 0.8y_i^{t-1} + y_i$ |
| $f_3(y_i, t) = y_i^{t-1} + y_i^t + y_i^{t+1} + y_i^{t+2}$ |
| $f_4(y_i, t) = y_i^t + y_i^{t+1} + y_i^{t+2} + y_i^{t+3}$ |

**Table S2.** Effect of different signal processing steps on connectome inference. AUROC and AUPR are reported for small datasets.

| Filter | AUROC |  |  |  |  |  | AUPR |  |  |  |  |  |
| --- | --- | --- | --- | --- | --- | --- | --- | --- | --- | --- | --- | --- |
|  | small-1 | small-2 | small-3 | small-4 | small-5 | small-6 | small-1 | small-2 | small-3 | small-4 | small-5 | small-6 |
| Spikes<br>$x$ | 0.781 | 0.827 | 0.872 | 0.901 | 0.907 | 0.938 | 0.412 | 0.460 | 0.516 | 0.592 | 0.609 | 0.667 |
| Spikes<br>+ Binarization<br>$u(x)$ | 0.542 | 0.521 | 0.533 | 0.558 | 0.554 | 0.673 | 0.204 | 0.182 | 0.170 | 0.180 | 0.171 | 0.224 |
| Spikes<br>+ Thresholding<br>$g(x)$ | 0.795 | 0.839 | 0.879 | 0.903 | 0.910 | 0.933 | 0.480 | 0.545 | 0.604 | 0.671 | 0.689 | 0.733 |
| Spikes<br>+ Smoothing<br>$h(x)$ | 0.849 | 0.859 | 0.881 | 0.903 | 0.913 | 0.942 | 0.486 | 0.489 | 0.501 | 0.554 | 0.571 | 0.625 |
| **FARCI**<br>**$h(g(x))$** | **0.876** | **0.890** | **0.910** | **0.928** | **0.934** | **0.960** | **0.667** | **0.686** | **0.700** | **0.767** | **0.787** | **0.838** |

**Table S3.** Effect of different signal processing steps on connectome inference. AUROC and AUPR are reported for normal datasets.

| Filter | AUROC |  |  |  | AUPR |  |  |  |
| --- | --- | --- | --- | --- | --- | --- | --- | --- |
|  | normal-1 | normal-2 | normal-3 | normal-4 | normal-1 | normal-2 | normal-3 | normal-4 |
| Spikes<br>$x$ | 0.891 | 0.894 | 0.892 | 0.889 | 0.334 | 0.337 | 0.342 | 0.329 |
| Spikes<br>+ Binarization<br>$u(x)$ | 0.656 | 0.655 | 0.655 | 0.643 | 0.042 | 0.043 | 0.044 | 0.040 |
| Spikes<br>+ Thresholding<br>$g(x)$ | 0.878 | 0.885 | 0.884 | 0.880 | 0.395 | 0.413 | 0.422 | 0.403 |
| Spikes<br>+ Smoothing<br>$h(x)$ | 0.910 | 0.909 | 0.910 | 0.906 | 0.336 | 0.327 | 0.332 | 0.327 |
| **FARCI**<br>**$h(g(x))$** | **0.907** | **0.909** | **0.910** | **0.905** | **0.490** | **0.491** | **0.495** | **0.488** |

**Table S4.** Effect of different signal processing steps on connectome inference.

| Filter | AUROC |  |  |  |  |  | AUPRC |  |  |  |  |  |
| --- | --- | --- | --- | --- | --- | --- | --- | --- | --- | --- | --- | --- |
|  | normal-3<br>(highrate) | normal-4<br>(lownoise) | highcc | lowcc | highcon | lowcon | normal-3<br>(highrate) | normal-4<br>(lownoise) | highcc | lowcc | highcon | lowcon |
| Spikes<br>$x$ | 0.937 | 0.919 | 0.907 | 0.870 | 0.867 | 0.930 | 0.409 | 0.366 | 0.416 | 0.233 | 0.321 | 0.334 |
| Spikes<br>+ Binarization<br>$u(x)$ | 0.742 | 0.625 | 0.707 | 0.569 | 0.579 | 0.660 | 0.091 | 0.039 | 0.054 | 0.028 | 0.046 | 0.044 |
| Spikes<br>+ Thresholding<br>$g(x)$ | 0.935 | 0.914 | 0.900 | 0.859 | 0.835 | 0.927 | 0.533 | 0.507 | 0.509 | 0.271 | 0.298 | 0.407 |
| Spikes<br>+ Smoothing<br>$h(x)$ | 0.945 | 0.913 | 0.912 | 0.888 | 0.894 | 0.949 | 0.405 | 0.337 | 0.392 | 0.255 | 0.334 | 0.350 |
| **FARCI**<br>**$h(g(x))$** | **0.953** | **0.915** | **0.916** | **0.892** | **0.875** | **0.956** | **0.602** | **0.532** | **0.575** | **0.380** | **0.368** | **0.525** |

**Table S5.** Comparison of FARCI performance with the winner algorithm of the Neural Connectomics Challenge on all the datasets provided in Neural Connectomics Challenge.

| Networks | AUROC |  | AUPRC |  |
| --- | --- | --- | --- | --- |
|  | FARCI | Sutera et al.<br>(2015) | FARCI | Sutera et al.<br>(2015) |
| small1 | 0.876 | 0.720 | 0.667 | 0.401 |
| small2 | 0.890 | 0.791 | 0.686 | 0.490 |
| small3 | 0.910 | 0.859 | 0.700 | 0.598 |
| small4 | 0.928 | 0.895 | 0.767 | 0.717 |
| small5 | 0.934 | 0.919 | 0.787 | 0.751 |
| small6 | 0.960 | 0.960 | 0.838 | 0.845 |
| normal1 | 0.907 | 0.943 | 0.490 | 0.403 |
| normal2 | 0.909 | 0.942 | 0.491 | 0.404 |
| normal3 | 0.910 | 0.942 | 0.495 | 0.398 |
| normal4 | 0.905 | 0.939 | 0.488 | 0.388 |
| n3-highrate | 0.953 | 0.961 | 0.602 | 0.452 |
| n4-lownoise | 0.915 | 0.941 | 0.532 | 0.432 |
| highcc | 0.916 | 0.946 | 0.575 | 0.486 |
| lowcc | 0.892 | 0.925 | 0.380 | 0.260 |
| highcon | 0.875 | 0.944 | 0.368 | 0.413 |
| lowcon | 0.956 | 0.955 | 0.525 | 0.334 |

